# Fabrication and delivery of mechano-actived microcapsules containing osteogenic factors in a large animal model of osteochondral injury

**DOI:** 10.1101/2021.09.24.461696

**Authors:** Hannah M. Zlotnick, Ryan C. Locke, Sanjana Hemdev, Brendan D. Stoeckl, Sachin Gupta, Ana P. Peredo, David R. Steinberg, James L. Carey, Daeyeon Lee, George R. Dodge, Robert L. Mauck

**Affiliations:** Department of Bioengineering, School of Engineering and Applied Sciences, University of Pennsylvania, Philadelphia, PA, USA; McKay Orthopaedic Research Laboratory, Department of Orthopaedic Surgery, Perelman School of Medicine, University of Pennsylvania, Philadelphia, PA, USA; Translational Musculoskeletal Research Center, Corporal Michael J Crescenz VA Medical Center, Philadelphia, PA, USA; Department of Biotechnology, School of Engineering and Applied Sciences, University of Pennsylvania, Philadelphia, PA, USA; Department of Chemical and Biomolecular Engineering, School of Engineering and Applied Sciences, University of Pennsylvania, Philadelphia, PA, USA

**Keywords:** osteochondral interface, bone, articular cartilage, controlled release, animal model, microcapsule

## Abstract

Chondral and osteochondral repair strategies are limited by adverse bony changes that occur after injury. Bone resorption can cause entire scaffolds, engineered tissues, or even endogenous repair tissues to subside below the cartilage surface. To address this translational issue, we fabricated poly(D,L-lactide-co-glycolide) (PLGA) microcapsules containing the pro-osteogenic agents triiodothyronine and ß-glycerophosphate, and delivered these microcapsules in a large animal model of osteochondral injury to preserve bone structure. We demonstrate that developed microcapsules ruptured *in vitro* under increasing mechanical loads, and readily sink within a liquid solution, allowing for gravity-based positioning onto the osteochondral surface. In a large animal, these mechano-active microcapsules (MAMCs) were assessed through two different delivery strategies. Intra-articular injection of control MAMCs enabled fluorescent quantification of MAMC rupture and cargo release in a synovial joint setting over time in vivo. This joint-wide injection also confirmed that the MAMCs do not elicit an inflammatory response. In the contralateral hindlimbs, chondral defects were created, MAMCs were locally administered, and nanofracture (Nfx), a clinically utilized method to promote cartilage repair, was performed. The NFx holes enabled marrow-derived stromal cells to enter the defect area and served as repeatable bone injury sites to monitor over time. Animals were evaluated 1 and 2 weeks after injection and surgery. Analysis of injected MAMCs showed that bioactive cargo was released in a controlled fashion over 2 weeks. A bone fluorochrome label injected at the time of surgery displayed maintenance of mineral labeling in the therapeutic group, but resorption in both control groups. Alkaline phosphatase (AP) staining at the osteochondral interface revealed higher AP activity in defects treated with therapeutic MAMCs. Overall, this study establishes a new micro-fluidically generated delivery platform that releases therapeutic factors in an articulating joint, and reduces this to practice in the delivery of therapeutics that preserve bone structure after osteochondral injury.

## 1. Introduction

Tissue interfaces are complex and notoriously challenging to engineer and regenerate. These boundaries are defined by continuous transitions in the cellular and extracellular matrix (ECM) components that define each tissue [1–3]. At the osteochondral interface, articular chondrocytes, hypertrophic chondrocytes, and osteoblasts are present in a graded manner [4]. Accordingly, from cartilage to bone there is an increase in mineralized ECM and decrease in collagen type II and proteoglycans [5]. Not only do these cellular and ECM components vary in concentration, but they also have markedly different organizations throughout the osteochondral unit. Notably, collagen fibers rotate from parallel at the cartilage surface to perpendicular at the osteochondral junction [6]. All together, these compositional gradients and organizational differences contribute to the mechanical strength of the osteochondral interface. This interface is severely disrupted in cases of cartilage injury [7–9], which are often accompanied by adverse bone changes (i.e., resorption, sclerosis, edema). Furthermore, in severe joint-wide disease that is sustained over long periods of time, subchondral bone changes may also further cartilage degeneration. Therefore, it is important to consider the entire osteochondral unit in the pursuit of an optimal repair strategy.

Biomimicry may enhance engineered tissue function and durability, and this has motivated considerable research in the field of gradient biofabrication. Numerous fabrication techniques have been explored to recreate chondral and osteochondral gradients. Remote fields (i.e., magnetic, gravitational, etc.) can be used to non-invasively pattern cells, and bioactive factors in a range of materials [10–13]. Additionally, three-dimensional (3D) weaving [14], electrospinning techniques [15,16], and custom molds can be used to provide different material stiffness cues to cells throughout the depth of engineered tissues [17]. More recently, gradient biofabrication via 3D printing has emerged a versatile approach [18–20], wherein one can easily position cells, bioactive factors, and/or material cues.

Despite this tremendous progress in the field of gradient biofabrication, clinical translation has been hindered by changes to the native bone structure that emerge with disease and at the time of material implantation. Bone resorption has been observed in rabbit [21], ovine [22], porcine [23,24], and equine models of cartilage and osteochondral repair [25,26], causing unwanted subsidence of both scaffold-based, engineered tissue and endogenous repair tissue. Ideally, the repair tissue would remain level with nearby articular cartilage to restore the articulating surface.

Here, we drew inspiration from the field of gradient biofabrication to deliver therapeutic microcapsules *in situ* to the osteochondral interface, with the goal of preserving and promoting subchondral bone formation following an osteochondral injury in a large animal. Specifically, we produced poly(D,L-lactide-co-glycolide) (PLGA) microcapsules with thick shells to enable *in situ* density-based positioning of these delivery vehicles to the osteochondral interface. The thick capsular shell was also used to control the rate of hydrolytic degradation within the synovial environment of the knee joint, making mechanical force the primary rupture mechanism. This mechanical force-based rupture profile was characterized in control and therapeutic microcapsules. The therapeutic mechano-activated microcapsules, or MAMCs [27–29], were fabricated to contain the bioactive therapeutics triiodothyronine (T3) and ß-glycerophosphate to stimulate stromal cell hypertrophy and matrix mineralization, respectively.

This study had two primary goals. The first goal was to assess progressive MAMC rupture over time in an *in vivo* articulating joint setting, and the impact of MAMC delivery on overall joint health. To do this, the left hindlimbs of six Yucatan minipigs received an intra-articular injection of control MAMCs. The second goal of this study was to assess the early impact of the therapeutic MAMCs containing agents that promote mesenchymal progenitor cell hypertrophy and biomineralization on bone repair and preservation in a surgical defect model. To do this, full thickness chondral defects were created in the right operative hindlimbs. Subsequently, MAMCs were locally administered to the osteochondral interface, and two nanofracture holes (1.2 mm diameter x 8 mm depth) were created [30], enabling stromal cells to enter the chondral defect. The nanofracture holes also served as a repeatable bone injury, and site to assess bone repair. Animals were evaluated 1 and 2 weeks after intra-articular injection and surgery. Overall, we show here that *in situ* delivery of MAMCs to the osteochondral interface presents a novel approach to locally apply therapeutics in a mechano-responsive manner with translationally relevant outcomes that potentially preserves bone structure after cartilage repair surgery.

## 2. Materials and methods

### 2.1 Materials

Penicillin/streptomycin/fungizone (PSF), phosphate buffered saline (PBS), Dulbecco’s Modified Eagle Medium (DMEM) (11965118), albumin from bovine serum Alexa Fluor™ 488 conjugate (Alexa Fluor™ 488 conjugated BSA) (A13100), chloroform (C298), TO-PRO-3 Iodide (T3605), Hoechst 33342 (H3570), and glacial acetic acid (A38) were purchased form Thermo Fisher Scientific (Waltham, MA). Bovine serum albumin (BSA) (A7906), T3 (T6397), ß-glycerophosphate disodium salt hydrate (G9422), Nile Red (N3013), poly(vinyl alcohol) (PVA) (P8136), sodium chloride (NaCl) (S5886), xylenol orange tetrasodium salt (398187), polyvinylpyrrolidone (PVP) (P5288), sucrose (S7903), sodium acetate anhydrous (W302406), and L-(+)-tartaric acid (251380) were purchased from Sigma Aldrich (St. Louis, MO). Transforming growth factor-ß3 (TGF-ß3) (8420-B3), and fetal bovine serum (FBS) (S11150) were purchased from R&D Systems (Minneapolis, MN). 85:15 lactide:glycolide PLGA (B6006-1) copolymer was purchased from Lactel (Birmingham, AL).

### 2.2 Pellet fabrication and culture

Mesenchymal stromal cells (MSCs) were isolated from juvenile bovine stifle joints (3 months old, Research 87, Marlborough, MA) as previously described [31]. MSCs were pooled from three donors, and were expanded in a basal media consisting of high glucose DMEM supplemented with 10% FBS and 1% PSF. At passage 2, MSCs were trypsinized and washed with PBS. MSC pellets (200,000 cells/pellet, *n* = 32 pellets) were formed in polypropylene V-bottom 96-well plates by centrifugation at 300 g for 5 min. All pellets were chondrogenically differentiated in chemically defined media (CM) with high-glucose DMEM, 1% PSF, dexamethasone (0.1 mM), sodium pyruvate (100 mg mL^-1^), L-proline (40 mg mL^-1^), ascorbate 2-phosphate (50 mg mL^-1^), insulin (6.25 mg mL^-1^), transferrin (6.25 mg mL^-1^), selenous acid (6.25 ng mL^-1^), BSA (1.25 mg mL^-1^), linoleic acid (5.35 mg mL^-1^), and TGF-ß3 (10 ng mL^-1^) [32,33]. After a 2-week pre-differentiation period, the pellets were washed with PBS, and the media was switched to CM without TGF-ß3 for the rest of the study. A single dose of T3 (10 nM) and ß-glycerophosphate disodium salt hydrate (10 mM), small molecules commonly added to osteogenic media, were administered to one half of the pellets [33]. The remaining pellets (control group, *n* = 16) did not receive any pro-osteogenic additives. For both the treatment and control groups, media was changed three times per week for 2 weeks. For determination of alkaline phosphatase (AP) activity, media was collected 1 week and 2 weeks post-treatment. A colorimetric AP assay kit (Abcam, Cambridge, UK, ab83369) was used to quantify AP content in the media per well.

### 2.3 Microcapsule fabrication and characterization

Microcapsules were fabricated using a glass capillary microfluidic device to generate water-in-oil-water double emulsions using three liquid phases [27–29,34]. Two batches of microcapsules were fabricated, referred to as control MAMCs and therapeutic MAMCs. The inner aqueous phase (pH 7.4) of both batches contained BSA (1 mg mL^-1^), and Alexa Fluor™ 488 conjugated BSA (0.01%). The inner aqueous phase of the therapeutic MAMCs also contained T3 (15.3 µM) and ß-glycerophosphate disodium salt hydrate (10 mM). The concentration of T3 was selected to deliver 1 ng of T3 in ∼3700 microcapsules (∼0.27 pg/microcapsule), matching the mass of T3 provided to a single MSC pellet in the *in vitro* studies. The middle and outer phases were the same for both MAMC batches. The middle phase consisted of 85:15 PLGA dissolved in chloroform with Nile Red (100 µg mL^-1^) for fluorescent visualization. The outer phase consisted of 2% PVA. Control and therapeutic MAMCs were separately collected in pH 12.08 phosphate buffered saline (PBS) with 0.1% BSA and NaCl (+0.95M). After collection, the MAMCs were left at 20°C for 72 h during which time the chloroform evaporated, and the shells solidified. The MAMCs were subsequently washed in PBS and stored at 4°C prior to use.

Microcapsule dimensions were assessed via confocal microscopy (Nikon A1R+, 10x magnification, mid-plane imaging) for each batch of MAMCs. The average outer diameter and shell thickness were calculated (ImageJ) from the mid-plane images (*n* = 50 MAMCs/batch). Additionally, the percentage of full MAMCs was determined from confocal images (4x magnification, mid-plane imaging). Briefly, the green channel was thresholded to quantify full MAMCs, and the total number of MAMCs, from which the percentage of full MAMCs (% full) was determined (*n* > 600 MAMCs/batch).

### 2.4 Mechanical assessment of microcapsules

To determine mechano-activation thresholds of the control and therapeutic MAMCs, a single layer of microcapsules (*∼*500) was placed between two glass coverslips and compressed at a controlled strain rate (0.5% s^−1^) to defined loads (1 and 5 N) using an Instron 5542. Unloaded (0 N) MAMCs served as negative controls. After application of load, MAMCs were collected in PBS and incubated overnight at 20°C to enable complete diffusion of the inner contents (Alexa Fluor™ 488 conjugated BSA) out of ruptured capsules. After this incubation, MAMCs were imaged on a confocal microscope (4x magnification, mid-plane imaging). Images were analyzed to quantify the % full after each load. The percentage of full MAMCs after load was normalized to the original % full level for each batch to account for differences in initial baseline levels.

### 2.5 Gravity-based positioning of microcapsules

To mimic the orthotopic *in vivo* delivery of the microcapsules, control microcapsules were injected into a PBS solution and serially imaged to visualize the timescale of settling. In vivo, the intent was for microcapsules to settle to the osteochondral interface. In this *in vitro* model, the microcapsules settled on the bottom surface of the vial. For this test, microcapsules (∼3700) were injected into saline (2.5 cm height) and allowed to sink over the course of 15 s.

### 2.6 Animal procedures and euthanasia

All animal studies were performed under an Institutional Animal Care and Use Committee-approved protocol at the University of Pennsylvania. Six skeletally mature (age 12 months at the beginning of the study) male Yucatan minipigs (Sinclair Bioresources, Auxvasse, MO, USA) were used for this study (figure 1(a)). Under a single anesthetic induction, two injections and one surgical procedure, as previously described [35], were performed.

**Figure 1.**
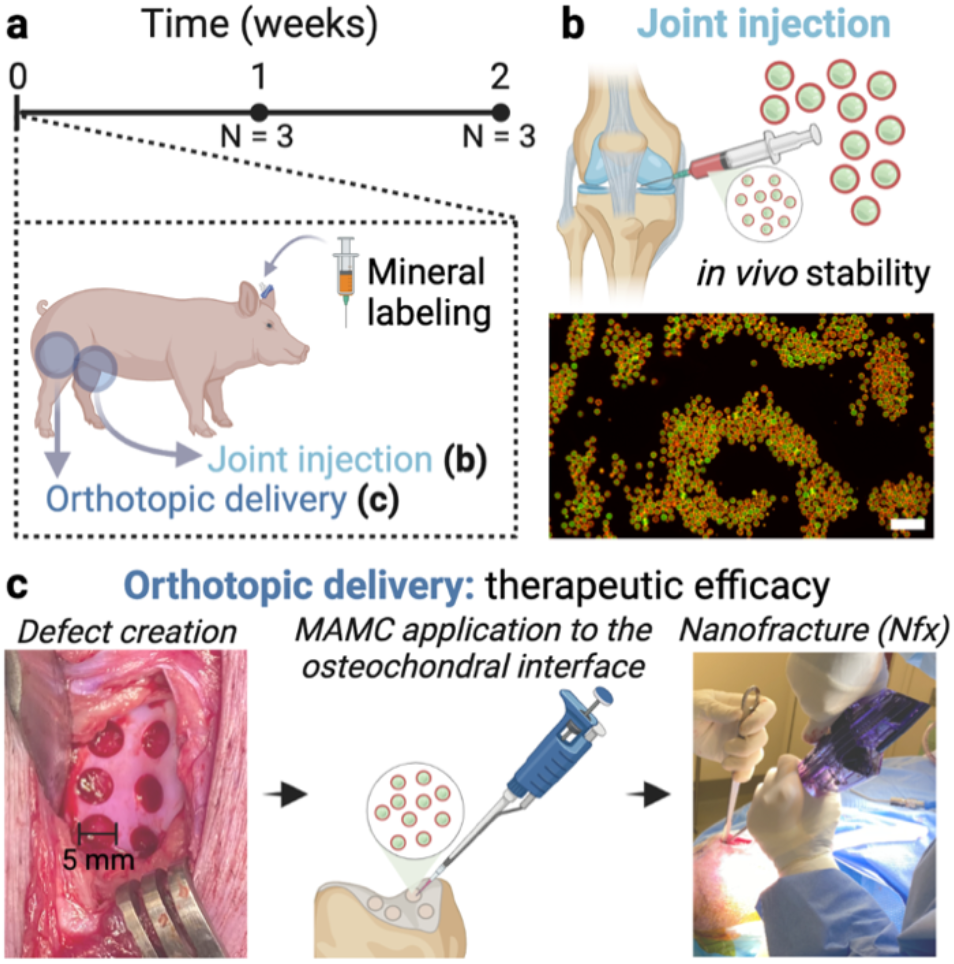
Microcapsule stability and therapeutic efficacy in a large animal model. (a) Yucatan minipigs were sacrificed 1 (N = 3 animals) and 2 (N = 3 animals) weeks after surgery/injection. A bone fluorochrome label was injected at time 0. (b) Microcapsules were intra-articularly injected in all left hindlimbs. Scale bar = 200 µm. (c) Microcapsules were delivered to the osteochondral interface to preserve bone after nanofracture in all right hindlimbs.

First, an ear vein catheter was placed to allow the perfusion of a bone fluorochrome label. Xylenol orange (90 mg kg^-1^) was injected over 1 h through the ear catheter [30,36]. The xylenol orange label was dissolved in 100 mL saline and sterile filtered prior to injection. Post-injection the catheter was flushed with 10 mL saline and removed.

After beginning the fluorochrome injection, the left hindlimb was sterilely prepared with betadine. An anterolateral approach was taken to inject control microcapsules (50,000 MAMCs per injection) into the knee joint (figure 1(b)). A 21-gauge 1½ inch needle was positioned perpendicular to the skin with the tip of the needle directed at a 45° angle into the center of the knee. Sterile saline was first injected to ensure proper placement of the needle. Next, a unilateral stifle joint procedure was performed on the right hindlimb (figure 1(c)) [30,35]. For this, a medial parapatellar skin incision was used to expose the trochlea. Six full-thickness chondral defects were created in the trochlear groove using a 5 mm biopsy punch and surgical curette. Care was taken to preserve the subchondral plate. Following the creation of cartilage defects, MAMCs were applied. Within each animal, two defects were left empty, two defects received control MAMCs, and two defects received therapeutic MAMCs. MAMCs (*n* = 3700) were suspended in saline (20 µL), and sterilely pipetted into the respective defects. The microcapsules were allowed to settle for 1 min. Finally, all defects received 2 nanofracture (Nfx) holes using the 1.2 mm diameter SmartShot^®^ marrow access device (Marrow Access Technologies, Minnetonka, MN, USA). This device created repeatable holes 8 mm into the subchondral bone. Experimental groups included: Nfx, Nfx + control MAMCs, and Nfx + therapeutic MAMCs. These groups were positionally randomized in each animal (*n* = 6/group/time point). Sample size was determined based on an *a priori* power analysis using previous data in this animal model [30]. All animals were ambulating within 2 h post-surgery. Animals were euthanized 1 (*N* = 3) and 2 (*N* = 3) weeks after surgery. The left (joint injection) and right (orthotopic delivery) hindlimbs were collected for separate analyses.

### 2.7 Quantification of intact microcapsules after joint injection

Three cadaveric Yucatan minipig hindlimbs (Sierra Medical, Whittier, CA, USA) were used to assess MAMC % full immediately after injection into a knee. The cadaveric limbs were injected with 50,000 MAMCs using a 21-gauge 1 ½ inch needle, and immediately dissected for analysis. The left hindlimbs from the primary animal study were concurrently dissected. The isolated tissues were imaged using an upright microscope (Zeiss AxioZoom.V16). Fluorescent images were captured to quantify MAMC % full over time in an articulating joint (0, 1, and 2 weeks).

### 2.8 Paraffin-embedded histology of synovium

After imaging the synovium to locate microcapsules, synovial tissues were isolated and fixed in 4% paraformaldehyde, embedded in paraffin, and sectioned (7 µm) using a paraffin microtome (RM2245, Leica). Sections were stained with haematoxylin and eosin (H&E) and imaged using a Zeiss AxioScan.Z1 digital slide scanner.

### 2.9 Micro-computed tomography and analysis

The right operative hindlimbs were dissected to isolate the femoral trochleae. A hand saw was used to isolate 1 mm^3^ osteochondral units, each containing a chondral defect (*n* = 36 total defects) [23]. The osteochondral units were immediately imaged by micro-computed tomography (µCT; Scanco µCT45, Scanco Medical). Scans were conducted utilizing the following parameters: 1,500 projections, 600 ms x 1 exposure/projection, voltage 55 kVp, current 145 µA, isotropic voxel size 10.4 µm. The µCT files were opened in Dragonfly software, Version 2020.2 for Windows, (Object Research Systems (ORS) Inc, Montreal, Canada) to acquire snapshots at the mid-plane of each defect. Subsequently, the Dragonfly software was used to calculate the bone volume/total volume within a cylindrical (5 mm diameter x 0.5 mm thickness) region of interest for each defect. The region of interest was aligned with the top and periphery of the defects.

### 2.10 Cryohistology of osteochondral units

Osteochondral units were fixed immediately after µCT scanning for 24 h in 10% neutral buffered formalin. After fixing, samples were infiltrated for 48 h with a 10 % sucrose and 2% PVP solution. Then, samples were embedded in OCT compound and sectioned (18 µm/section) undecalcified using cryofilm until reaching the midplane of the defect using a motorized cryostat (CM1950, Leica) [37]. Tape-stabilized, frozen sections were subjected to two rounds of imaging using a Zeiss AxioScan.Z1 digital slide scanner, including (i) fluorochrome label with TO-PRO-3 Iodide nuclear counterstain, and (ii) AP fluorescent staining with Hoechst 3342 nuclear counterstain [38]. Before AP staining, sections were decalcified for 1 h in 0.92% sodium acetate anhydrous, 1.14% L-(+)-tartaric acid, and 1% glacial acetic acid pH 4.1-4.3. AP staining occurred according to the manufacturer’s protocol (Vector Laboratories, Burlingame, CA, USA, SK-5300).

### 2.11 Statistical analyses

All quantitative data were analyzed using GraphPad Prism (version 8.4.3 for MacOS, GraphPad Software). For the assessment of microcapsule rupture over time *in vivo*, the xylenol orange quantification, and AP activity *in vivo* at a given time point a one-way analysis of variance (ANOVA) was performed with *post-hoc* Tukey’s corrections for multiple comparisons. For the in vitro pellet study, MAMC compression test, and bone resorption data, a two-way ANOVA was performed with post-hoc Tukey’s corrections for multiple comparisons. Significance was set at *p* < 0.05 with *, **, ***, **** indicating *p* <z 0.05, 0.01, 0.001, or 0.0001, respectively. All data are represented as mean ± standard deviation. All figures were created with BioRender.com.

## 3. Results and discussion

### 3.1 Osteogenic induction of MSC pellets

T3 and ß-Glycerophosphate are attractive candidates to include in a targeted delivery system for osteochondral preservation and repair. Thyroid hormones, including both T3 and thyroxine (T4), induce hypertrophy of mesenchymal stromal cells and chondrocytes [39,40], and ß-glycerophosphate provides raw materials to support the formation of mineralized extracellular matrix [41]. Both factors are cost-effective and soluble at high concentrations, allowing therapeutic dosages to be encapsulated in micron-scale drug delivery carriers. Here, T3 and ß-glycerophosphate were first administered to MSC pellets to assess whether a single delivery event of these factors could enhance alkaline phosphatase activity, an early indicator of hypertrophy and eventual osteogenesis. While it is well established that prolonged exposure to T3 and ß-glycerophosphate promotes cell hypertrophy and bone formation [33,42–44], it was not yet known whether a single dose could elicit such a response. Marrow-derived MSC pellets were first pre-differentiated in chondrogenic media for 2 weeks before initiating hypertrophy through the delivery of one dose of T3 and ß-glycerophosphate, and the removal of TGF-ß3 (figure 2(a)). Culture media was collected both 1 and 2 weeks after delivering the hypertrophic factors. An alkaline phosphatase assay on the collected media revealed stark differences between the control group (no treatment), and pellets that received the hypertrophic factors (figure 2(b)). As expected, there was an increase in AP activity in the media 1 week after administration of the hypertrophic factors. Surprisingly, AP activity was over 3 times higher at the 2-week time point compared to the 1-week time point within the therapeutic group, showing prolonged effect of a single delivery event. These data supported the encapsulation of T3 and ß-glycerophosphate within mechano-active microcapsules, to control the localization and release timing of these pro-hypertrophic and pro-osteogenic factors *in vivo*.

**Figure 2.**
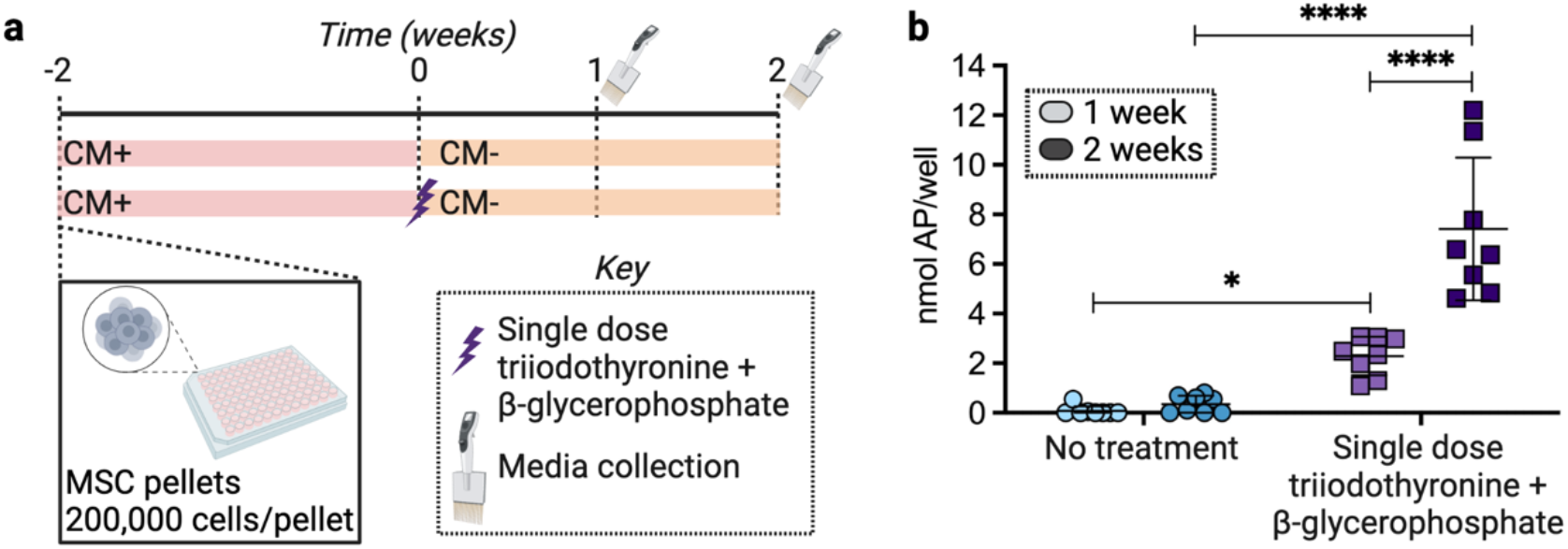
Osteogenic induction of mesenchymal stromal cell (MSC) pellets. (a) Experimental overview of pellet study. Pellets were first cultured in chondrogenic media with 10 ng mL^-1^ TGF-β3, and then switched to chemically defined media without TGF-β3. The treatment group received a single dose of osteogenic factors at time 0. (b) Alkaline phosphatase (AP) present in the collected culture media 1 week and 2 weeks after treatment (mean ± standard deviation, n = 8 pellets/condition/time point, * p < 0.05. **** p < 0.0001).

### 3.2 Fabrication of mechano-active microcapsules

To establish a clinically relevant drug delivery system, production scalability is an important consideration. Here, glass capillary microfluidic devices were used to enable high-throughput fabrication of both control and therapeutic microcapsules (figure 3(a)). Within 2 h, > 2 million MAMCs were produced. These devices had three inlet ports for the outer (PVA), middle (PLGA + Nile Red), and inner (Alexa Fluor™ 488 conjugated BSA ± therapeutics) phases, respectively. Both control and therapeutic MAMCs were fabricated, with the only difference being that the inner phase of the therapeutic MAMCs contained T3 and ß-glycerophosphate. While other bioactive factors have previously been included in double emulsions [27,28], this was the first time that T3 and ß-glycerophosphate were encapsulated in such microcapsules. Inclusion of these hypertrophic additives did not disrupt the fabrication process.

**Figure 3.**
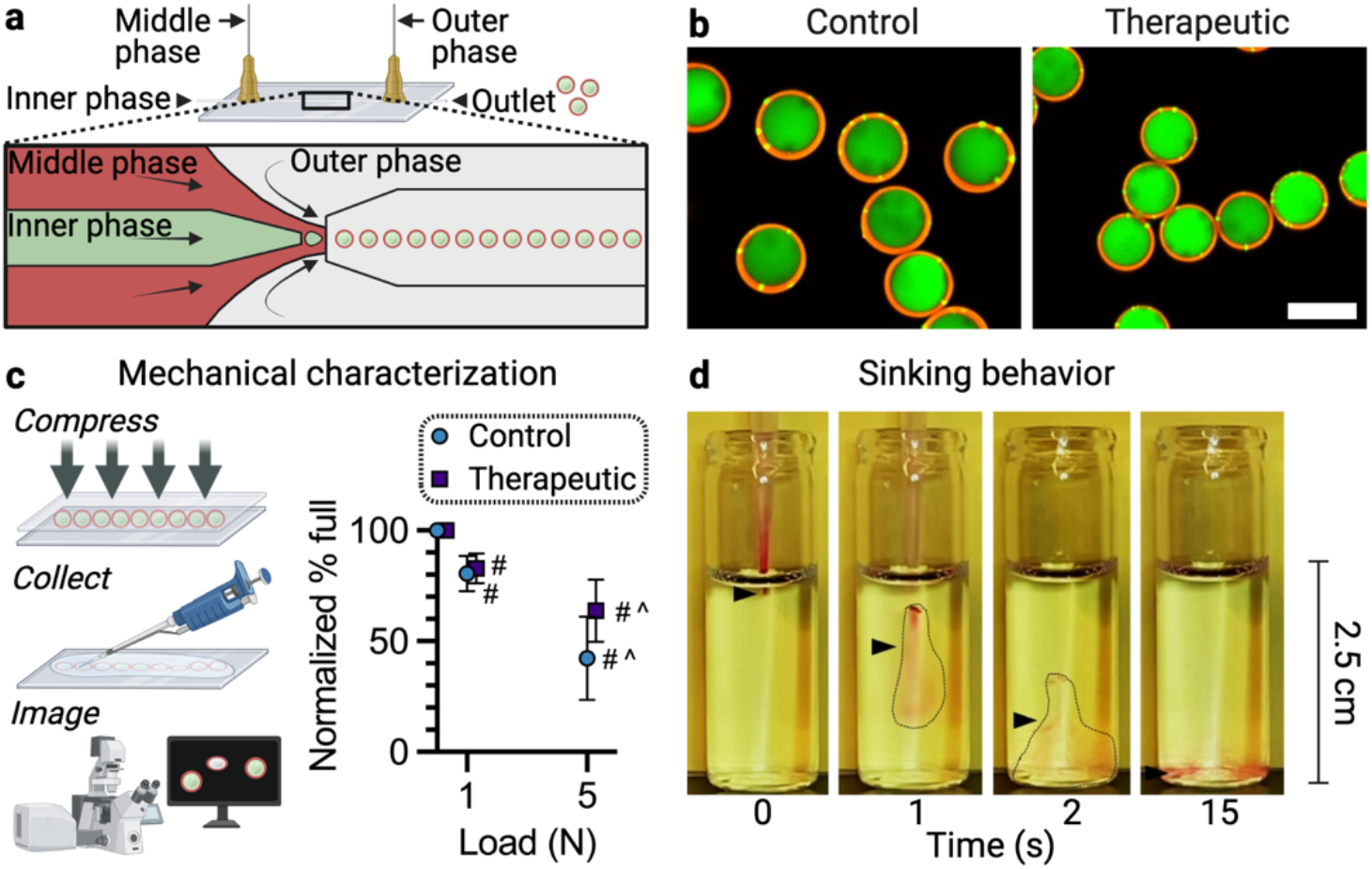
Fabrication of mechanically activated microcapsules (MAMCs) containing osteogenic factors. (a) Microfluidic fabrication of MAMCs. (b) Mid-plane confocal images of control and therapeutic MAMCs. Scale bar = 50 µm. (c) Mechanical characterization of MAMCs under direct static compression (mean ± standard deviation, n = 500 MAMCs/type/run, 5 runs MAMC type, # p < 0.05 vs. unloaded MAMCs, ^ p < 0.05 vs. MAMCs of the same type at 1N). (d) Sinking behavior of the MAMCs within a saline solution to mimic in vivo delivery.

Nile Red and Alexa Fluor™ 488 conjugated BSA allowed for visualization of the microcapsule shells, and inner contents, respectively, showing monodisperse double emulsions (figure 3(b)). The control MAMCs (D_control_: 48.56 µm ± 1.60 µm) had a slightly larger diameter than the therapeutic MAMCs (D_ther_: 39.23 µm ± 1.19 µm). Both batches of MAMCs had relatively thick PLGA shells (t_control_: 3.94 µm ± 1.07 µm, t_ther_: 2.31 µm ± 0.68 µm). These shell thickness measurements mirror a previous study that osmotically annealed PLGA microcapsules prior to shell hardening [28], as done in our study. In comparison, PLGA microcapsules collected in isosmotic solutions have shells that are 2 to 7 times thinner [27,29]. Thicker shells provide more resistance to hydrolytic degradation [28], which is desirable for the clinical translation in synovial joint spaces.

Both the control and therapeutic MAMCs progressively ruptured with increasing compressive loads (figure 3(c)), demonstrating their mechano-activation capacity. Despite differences in diameter and shell thickness, the control and therapeutic MAMCs had similar mechano-activation profiles. There were no differences in the percentage of full microcapsules between batches at a given load. This is likely because the control MAMCs with thicker shells (increasing load resistance), also had a larger diameter (decreasing load resistance).

Depending on the buoyant and gravitational forces at play on a given microcapsule, it can either float or sink in solution. For orthotopic delivery to the osteochondral interface, it was important to verify that microcapsules would rapidly sink within saline and percolate into the bone. Once the microcapsules settled, the saline carrier solution could leave the defect space. To estimate this settling time, we established a simplified model of the microcapsules’ descent (figure 3(d)). MAMCs (∼3700)—the same amount as in the orthotopic application— were injected into a saline solution. The MAMCs rapidly settled on the bottom surface of the vial within 15 s, a suitable timeframe for surgical application.

### 3.3 Stability of microcapsules in the synovial joint

To date, the stability of PLGA microcapsules has been assessed *in vitro* via prolonged incubation in synovial fluid, and *in vivo* via subcutaneous implantation in rats [28]. However, stability of these PLGA microcapsules has not previously been evaluated in a large animal synovial joint, which undergoes extensive loading with daily activity. Here, control MAMCs (50,000 within 1 mL saline) were injected into the left hindlimbs of six Yucatan minipigs, and the animals were evaluated 1- and 2-weeks post injection. Three additional cadaveric hindlimbs were similarly injected for a time zero assessment. Careful dissection of the knee joints post-injection revealed that the MAMCs were distributed throughout the joint space. In the limbs dissected immediately after injection, ∼95% of the MAMCs were full (figure 4(a, b)). This finding indicate that the injection alone did not rupture the MAMCs. After 1 week *in vivo*, ∼23% of the MAMCs remained full, and ∼10% of the MAMCs were full after 2 weeks *in vivo*. Therefore, PLGA microcapsules can be used to effectively extend drug residence time in a synovial joint. In this study, 75% of the cargo was delivered within the first week, 13% within the second week, with 10% remaining for delivery beyond the 2-week mark. Absent this sequestration within the MAMCs, the small molecule therapeutics used here would be expected to leave the joint within several hours [45].

**Figure 4.**
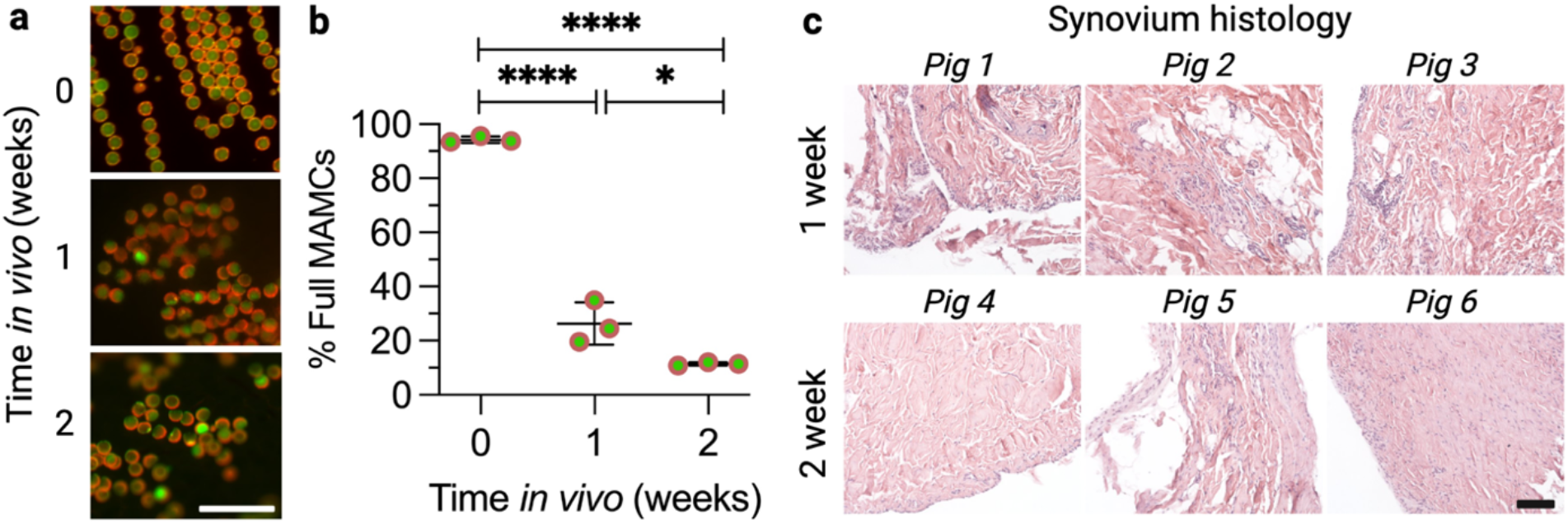
Mechano-activation of control microcapsules in the knee joint and impact on synovium over time in vivo. (a) Control microcapsules were imaged 0, 1, and 2 weeks after injection into the left hindlimbs of Yucatan minipigs (N = 3 animals/time point). Cadaveric Yucatan minipig hindlimbs were used for the time 0 assessment. Scale bar = 200 µm. (b) Percentage of microcapsules that retained their initial cargo as a function of time *in vivo* (mean ± standard deviation, N = 3 animals/time point, n = 100 microcapsules imaged/animal/time point, * p < 0.05, **** p < 0.0001). (c) Haematoxylin and Eosin (H&E) staining of synovium samples taken from injected hindlimbs. Each image represents a unique animal. Scale bar = 100 µm.

Synovium samples were taken from the suprapatellar region and processed for H&E staining to determine whether the PLGA microcapsules elicited any unwanted immune response (figure 4(c)). Representative images shown from each of the animals revealed mild hyperplasia, darker staining, and some vessels at the 1-week time point. However, by the 2-week time point, the synovium appeared lighter with a less cellular intimal lining, indicative of healthy tissue. Overall, the MAMCs had no apparent lasting impact on the synovium or other joint structures.

### 3.4 Micro-computed tomography analysis of defects

Bony abnormalities, including subchondral resorption, often complicate the long-term outcomes of chondral and osteochondral repair procedures, motivating localized treatment and preservation of subchondral bone [23,24,30,46,47]. Here, mechano-active microcapsules containing pro-osteogenic factors—T3 and ß-glycerophosphate—were delivered to the osteochondral interface to preserve subchondral bone after nanofracture. This study focused on the early bony changes that occurred 1- and 2-weeks post-surgery.

First, the osteochondral units were scanned via µCT (figure 5(a)). The bone volume/total volume was quantified for each sample (figure 5(b)). The 1-week values for all groups ranged from 4.3 – 4.8 with no significant differences between conditions (shaded region). The mean bone volume/total volume decreased from the 1-week to the 2-week time point across all groups, signifying bone loss, though the loss in bone was less pronounced in the therapeutic MAMC group. At this 2-week time point, there was considerable variability in this metric. Indeed, there appeared to be two subpopulations within the therapeutic group—the two lowest values, and the four higher values. It is possible that some of the defects better retained the microcapsules at the osteochondral interface, leading to a higher therapeutic efficacy in some defects.

**Figure 5.**
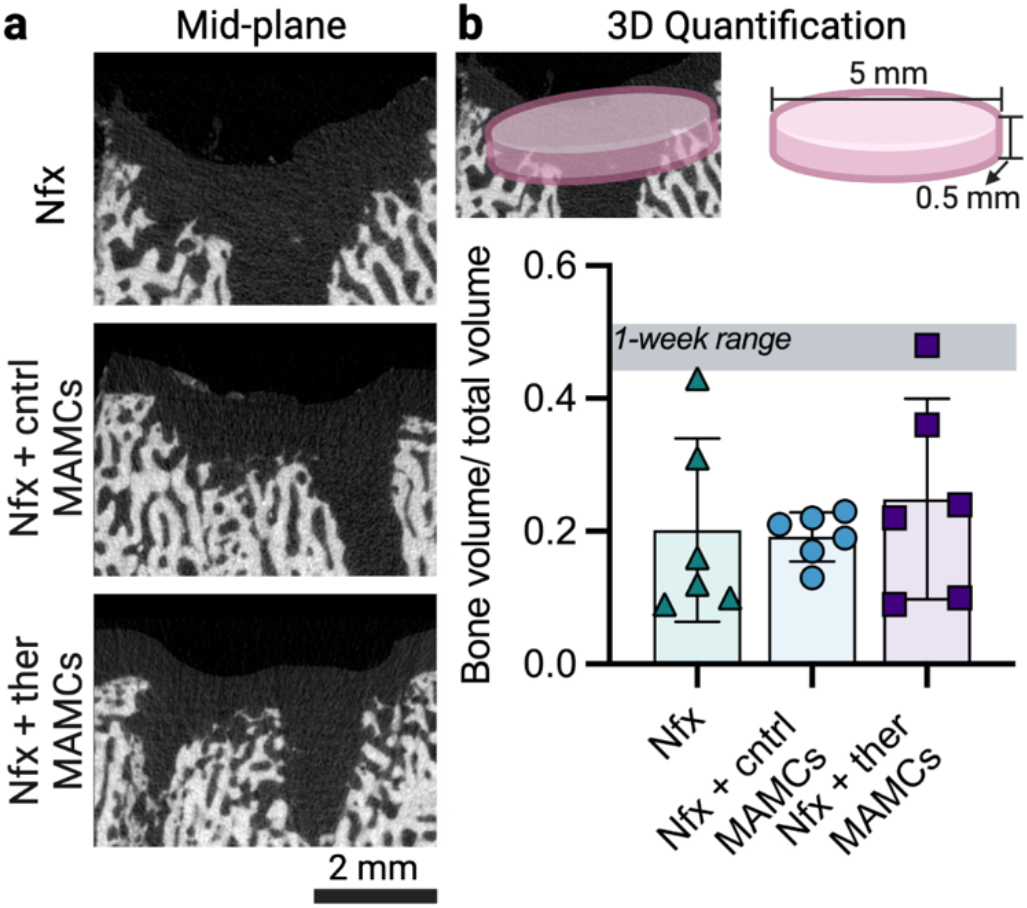
Bone volume/total volume following nanofracture (Nfx) and local mechano-active microcapsule (MAMC) delivery in a large animal. (a) Mid-plane image of the median defects for each of the groups at the 2-week time point, as assessed by µCT. (b) Schematic highlighting the cylindrical region of interest, and the volumetric quantification of bone volume/total volume 2 weeks post-surgery (mean ± standard deviation, n = 6 defects/group/time point, * p < 0.05). The shaded region represents the range of values 1-week post-surgery.

### 3.5 Bone mineral labeling with treatment

Fluorochrome labeling has been widely used in small animal studies (mouse, rat, rabbit, etc.), but has been less practiced in larger species (sheep, goat, pig, etc.) [48]. Here, we adapted a bone labeling protocol, previously described [30], to reduce the need for specialized personnel. Instead of implanting an indwelling jugular catheter, an ear vein catheter was used to intravenously inject xylenol orange at the time of surgery. There were no adverse effects of this procedure.

The bone mineral labeling demonstrated 1) the amount of newly mineralized bone (1-week time point), and 2) the disappearance or maintenance of this newly formed bone (2-week time point). Representative images show that the labeling lined the periphery of the nanofracture holes, or the site of ‘bone injury’ (figure 6(a)). There was little to no signal away from the nanofracture holes. For the quantification of these images, a smaller region of interest was established immediately adjacent to each nanofracture hole (figure 6(b)). Interestingly, within both the Nfx and Nfx + control MAMCs groups, there was a decline in labeling from the 1-week to the 2-week time point, signifying bone resorption. However, within the therapeutic group, xylenol orange labeling was maintained, highlighting improved subchondral bone preservation near the nanofracture holes. It is possible that some of the locally administered MAMCs fell into the nanofracture holes at the time of surgery, releasing their cargo. This would explain why we see a therapeutic benefit of the MAMCs in the area adjacent to the nanofracture holes.

**Figure 6.**
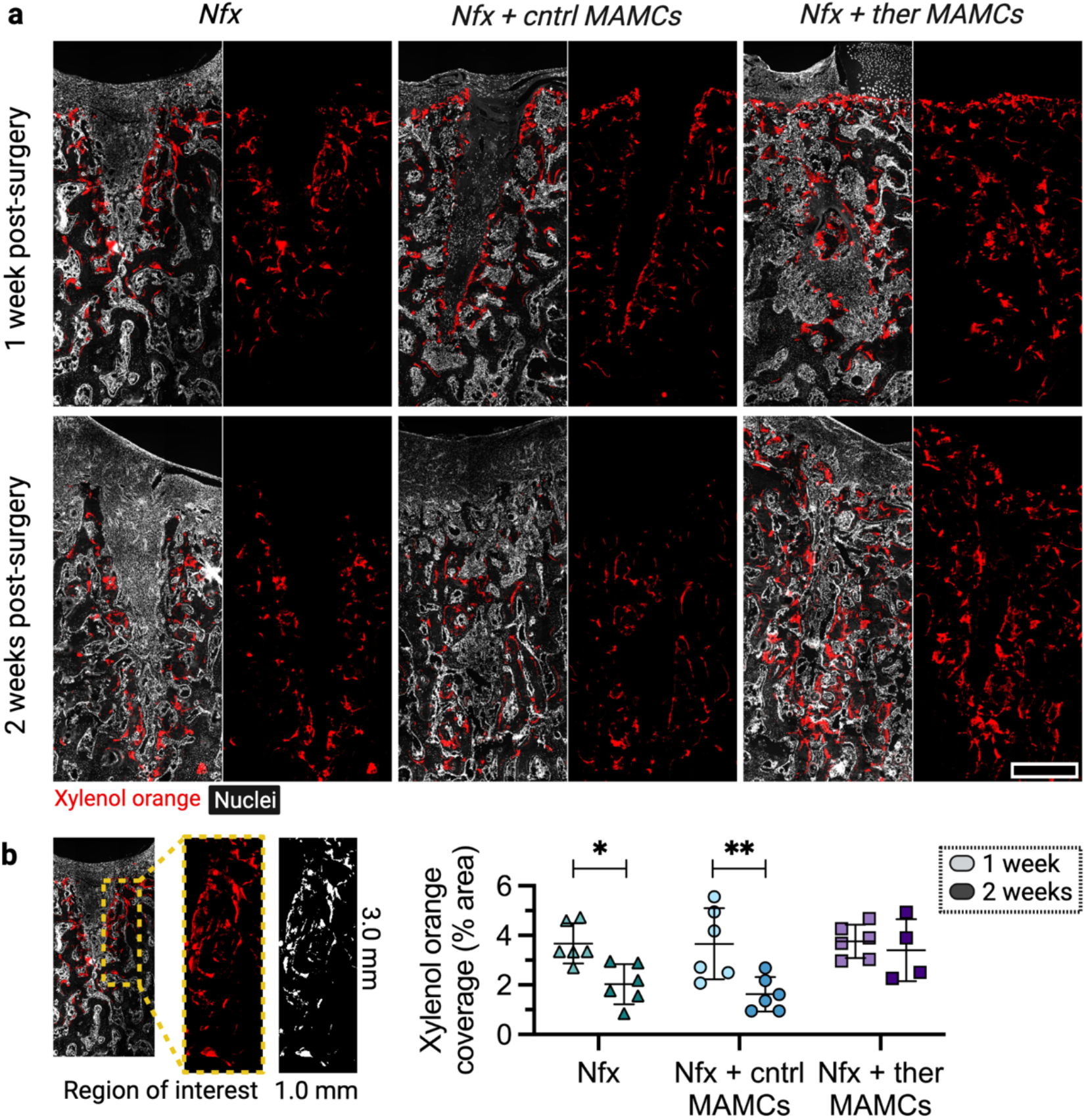
Therapeutic mechano-activated microcapsules (MAMCs) preserve bone mineral labeling adjacent to nanofracture (Nfx) holes. (a) Representative images from each of the groups 1 and 2 weeks after surgery. Xylenol orange was injected at the time of surgery. Red: xylenol orange, white: nuclei. Scale bar = 1.0 mm. (b) A region of interest (3 mm x 1 mm) was set adjacent to a Nfx hole. Xylenol orange coverage (% area) was quantified (mean ± standard deviation, n = 4-6 defects/group/time point, * p < 0.05, ** p < 0.01).

While we could not assess MAMC rupture over time in the operative limbs as we could in the contralateral limbs that received an intra-articular joint injection, the mineral labeling data suggest that, between the 1-week and 2-week endpoints, there was a therapeutic effect. This may be due to the gradual rupture of the MAMCs at the interface as the animals ambulated.

### 3.6 Alkaline phosphatase activity at the subchondral interface

AP staining of the defects was particularly striking at the osteochondral interface (figure 7(a)). A region of interest was set spanning the length of this interface for each sample. At both the 1- and 2-week time points, the defects treated with therapeutic MAMCs had elevated AP activity, as seen in the images (figure 7(b)) and image quantification (figure 7(c)). At the 2-week time point, the % area of the AP staining in the therapeutic group was two-fold higher than the Nfx group.

**Figure 7.**
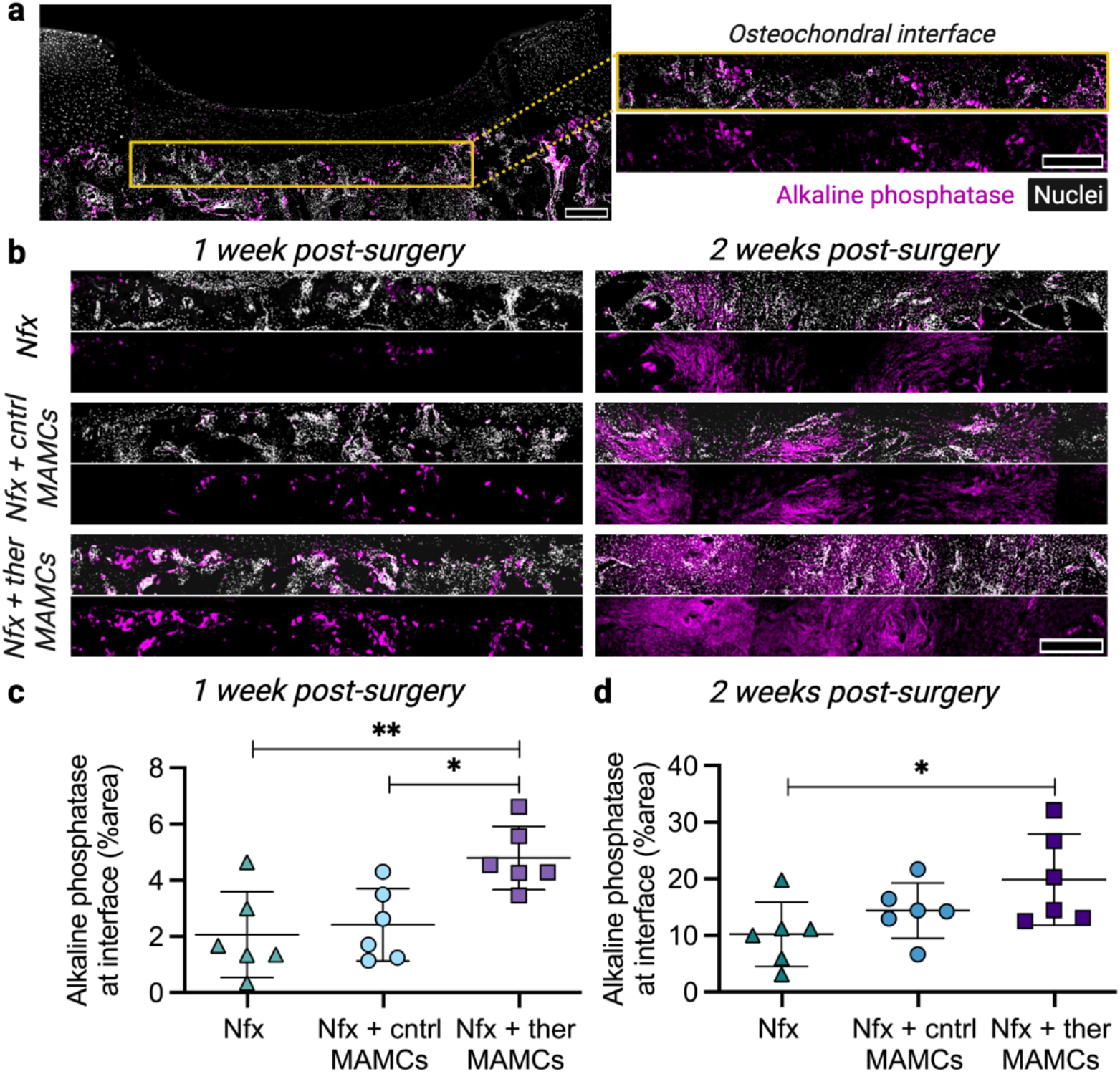
Therapeutic mechano-activated microcapsules (MAMCs) increase alkaline phosphatase activity at the osteochondral interface. (a) A region of interest (4 mm x 0.5 mm) was created at the osteochondral interface. Magenta: alkaline phosphatase, white: nuclei. Scale bars = 500 µm. (b) Representative images from each of the groups 1 week and 2 weeks after surgery. Nfx: nanofracture. Scale bar = 500 µm. (c, d) Quantification (% area) of alkaline phosphatase staining 1 and 2 weeks after surgery, respectively (mean ± standard deviation, n = 6 defects/group/time point, * p < 0.05, ** p < 0.01).

In accordance with the mineral labeling data, this data supports the prolonged effect of the bioactive therapeutics, beyond the 1-week time point. This is consistent with our *in vitro* pellet model, where we also saw an increase in AP two weeks after treating MSC pellets with a single dose of T3 and ß-glycerophosphate. The dosage of T3 and ß-glycerophosphate within the MAMCs was established from this initial pellet study. This suggests that ∼1 ng of T3 has a therapeutic impact on a ∼20 mm^2^ surface area of bone.

The mineral labeling and AP staining revealed a clear therapeutic benefit of the osteogenic MAMCs, despite the lack of significance in the µCT data. It is possible that with a greater animal number, we could identify outliers in each of our tested groups. While all large animal studies are limited by sample number, an efficient utilization of animals was demonstrated, and significant differences were identified between groups. A limitation of this study is that we were unable to assess MAMC retention in the defects themselves, given that repair tissue grew over the osteochondral interface (as intended in this standard cartilage repair procedure). MAMC retention in the defects could be further enhanced via hydrogel mediated MAMC delivery. In this study, we kept the orthotopic delivery method as translatable as possible, limiting the time of application and required materials. Future work will explore the long-term benefit of the pro-osteogenic microcapsules, and the inclusion of other bone promoting agents to further preserve subchondral bone after osteochondral injury, as well as target the overlying cartilage that forms in this approach.

## 4. Conclusion

Mechano-activated therapeutics are especially promising for dynamically loaded musculoskeletal tissues. Here, we were able to effectively encapsulate bioactive pro-osteogenic factors in PLGA microcapsules that rupture upon application of a physiological load. Upon injection into the knee joint, the control MAMCs remained full after delivery, and progressively ruptured over two weeks *in vivo*. The presence of MAMCs showed no obvious toxic or inflammatory response, motivating their further use in an orthotopic setting. In a first of its kind approach, therapeutic MAMCs containing pro-osteogenic factors were directly applied to the osteochondral interface to preserve bone structure after nanofracture, or bone ‘injury.’ These therapeutic MAMCs increased bone formation adjacent to the nanofracture holes, and at the osteochondral interface. Overall, this is the first assessment of the therapeutic potential of mechano-actived microcapsules in a large animal joint, and these findings demonstrate the promise of two different clinically relevant delivery approaches, intra-articular injection, and direct administration in a defect to improve joint health and direct tissue repair. Of translational relevance, this MAMC delivery platform permits the targeted application of therapeutics and ultimately, a new way to improve local tissue homeostasis and regeneration.

## 5. Acknowledgements

This work was supported by the Department of Veterans’ Affairs (I01 RX003375, IK6 RX003416), the NSF (CMMI: 15-48571), and the NIH/NIAMS (R01 AR071340, T32 AR007132, P30 AR069619, F31 AR077395). The authors would also like to thank Marrow Access Technologies for donating the SmartShot^®^ devices, and Beth Lemmon and the ULAR staff for their assistance with the animal surgeries and care. GRD is an employee of, and RLM and DL are co-founders of Mechano-Therapeutics, LLC.

